# Improved HLA-based prediction of coeliac disease identifies two novel HLA risk modifiers, DQ6.2 and DQ7.3

**DOI:** 10.1101/561308

**Authors:** Michael Erlichster, Justin Bedo, Efstratios Skafidas, Patrick Kwan, Adam Kowalczyk, Benjamin Goudey

## Abstract

**Purpose:** Human Leukocyte Antigen (HLA) testing is useful in the clinical work-up of coeliac disease (CD), with high negative but low positive predictive value. We construct a genomic risk score (GRS) using HLA risk loci to improve CD prediction and guide exclusion criteria.

**Methods:** Imputed HLA genotypes for five European CD case-control GWAS (n>15,000) were used to construct and validate an HLA based risk models (*HDQ*_15_). Conditioning on this score, we identified novel HLA interactions which modified CD risk, and integrated these novel alleles into a new risk score (*HDQ*_17_).

**Results:** A GRS from HLA risk allele genotypes yields performance equivalent to a state-of-the-art GRS *(GRS*_228_) using 228 single nucleotide polymorphisms (SNPs) and significantly improves upon all previous HLA based risk models. Conditioning on this model, we find two novel associations, HLA-DQ6.2 and HLA-DQ7.3, that interact significantly with HLA-DQ2.5 (p = 2.51 × 10^−9^, 1.99 × 10^−7^ for DQ6.2 and DQ7.3 respectively). These epistatic interactions yield the best performing risk score (*HDQ*_17_) which retains performance when implemented using 6 tag SNPs. Using the *HDQ*_17_ model, the positive predictive value of CD testing in high risk populations increases from 17.5% to 27.1% while maintaining a negative predictive value above 99%.

**Conclusion:** Our proposed HLA-based GRS achieves state-of-the-art risk prediction, helps elucidate further risk factors and improves HLA typing exclusionary criteria, which may reduce the number of patients requiring unnecessary endoscopies.

## Introduction

Coeliac disease (CD) is a chronic immune disease characterised by small intestine damage resulting from ingestion of gluten, the alcohol insoluble protein in wheat, barley and rye [1]. CD is a common disease, with a prevalence of 0.5–2% in Caucasian and Middle Eastern populations [2–4]. The current diagnostic gold-standard for CD is the demonstration of characteristic small intestinal inflammation and damage while on a gluten-containing diet [5]. Intestinal biopsies are obtained by upper gastrointestinal endoscopy, a resource intensive, invasive and inconvenient process [6]. While CD-specific serotyping of antibody markers is a strong positive predictor of disease, these tests are inaccurate in patients already on a gluten-free diet and patients with other conditions such as liver disease, inflammatory bowel disease and type-1 diabetes [7]. Alternative, non-invasive, risk stratification strategies are desired to reduce unnecessary endoscopies and improve the overall effectiveness of CD investigation [3].

One increasingly adopted strategy involves human leukocyte antigen (HLA) typing based on the exceptionally strong association of three major susceptibility alleles, HLA-DQA1*05, HLA-DQB1*02 and HLA-DQB1*03:02 with CD[8]. Over 99% of individuals with CD carry these risk alleles as part of the DQ2.5 (HLA-DQA1*05, HLA-DQB1*02), DQ2.2 (HLA-DQA1*02, HLA-DQB1*02), DQ7.5(HLA-DQA1*05 without HLA-DQB1*02) or DQ8 (HLA-DQB1*03:02) genotypes [9]. HLA genotyping, as currently used, has limited predictive value for CD due to the high population frequency of these susceptibility genotypes (30–60%)[10, 11]. Thus, while HLA typing alone cannot yield a diagnosis, the test is useful in selected clinical situations, such as assessing individuals already on a gluten-free diet[12]. In these scenarios, the strong negative predictive value of genetic testing can be used to confidently exclude a diagnosis of CD[8]. HLA genotyping is also valuable to help identify high-risk individuals, such as first-degree relatives of CD patients, where prevalence of disease is approximately 10% [8, 13].

While HLA typing is primarily used for exclusion, whereby those at ‘low’ risk (DQ7.5 or DQX) are excluded from diagnostic follow-up[6], there is evidence that this approach may not have captured the diversity in the relative risk of HLA genotypes. Firstly, recent clinical guidelines recommend further refinement of HLA genotype groupings into six categories based on observed relative risk, with the intent of more fine-grained risk estimates of high risk individuals [8]. Secondly, interactions between HLA haplotypes in CD, particularly the DQ2.2. and DQ7.5 haplotypes, were found to increase disease risk and better explain phenotypic variance[14, 15]. Thirdly, a logistic regression using a reduced set of HLA tag SNPs improved prediction of disease compared with coarse stratification, although this analysis did not account for DQ7.5 attributed risk [16]. These insights suggest that more nuanced stratification of risk HLA-DQ genotypes could be used to more effectively capture genetic risk.

To identify additional genomic variants that modulate CD risk, multiple groups have explored the use of loci outside of the HLA region by analysing genome-wide association studies (GWAS). Romanos *et al.* demonstrated that the addition of 57 non-HLA single nucleotide polymorphisms (SNPs) as part of a genomic risk score (GRS) could improve patient stratification over HLA genotype alone [17]. Abraham *et al.,* using GWAS data of five European populations, employed penalised regression models to develop a GRS using 228 SNPs capable of further improved patient stratification[16].

While these models represent the state-of-the-art for genomic prediction of CD, their role and the role of non-HLA genetic information in clinical practice is not yet well established, nor is this type of data routinely collected.

Here, we sought to determine whether the use of state-of-the-art statistical methods could improve prediction of CD with HLA genotypes using only HLA genotypes which are currently clinically collected. Using five European CD case-control GWAS datasets, we demonstrate that HLA-DQ genotype stratification has much greater predictive performance than previously attributed. By conditioning on this risk score, we identify two novel risk alleles, DQ6.2 and DQ7.3, in the UK2 cohort that show significant, replicating interaction effects with DQ2.5. Integrating these novel interactions into our risk score significantly improves risk prediction. Furthermore, we demonstrate that this model can be implemented using only six HLA tagging SNPs. Finally, we assess the impact of shifting the CD exclusionary criteria, demonstrating that it is possible to double the number of individuals excluded via genetic testing, with minimal impact on negative predictive value.

## Methods

### Genotype and Phenotype Data

Five European CD case and control datasets were used in this analysis from four populations: United Kingdom (UK1, UK2), Finland (FIN), Italy (IT) and the Netherlands (NL) (**Table 1**). Details regarding collection and quality control of these datasets have been described previously [18]. Previous analysis indicates population structure does not play a role in the predictive capacity of models built on these cohorts [16, 18].

**Table 1:** European CD GWAS cohorts used in this study.

| Population | SNP dataset | Post QC SNPs | Cases | Controls | Dataset Accession |
| --- | --- | --- | --- | --- | --- |
| UK1 | Illumina Hap300 | 295,453 | 737 | 2,596 | EGAD00010000292 |
| UK2 | Illumina Hap550 | 528,969 | 1,849 | 4,936 | EGAD00010000286 |
| FIN | Illumina Hap550 | 528,969 | 647 | 1,829 | EGAD00010000286 |
| NL | Illumina Hap550 | 528,969 | 803 | 846 | EGAD00010000286 |
| IT | Illumina Hap550 | 528,969 | 497 | 543 | EGAD00010000286 |
| Total |  |  | 4,533 | 10,750 |  |

### Imputation and grouping of HLA-DQA1 and HLA-DQB1 alleles

The R package HIBAG (HLA Genotype Imputation with Attribute Bagging) was used to impute 4-digit HLA-DQA1 and HLA-DQB1 genotypes for each sample [19]. Median posterior probability was 0.99 for imputations in the UK2, FIN, NL and IT populations and 0.90 for imputation in the reduced SNP UK1 dataset. Imputed HLA-DQA1 and HLA-DQB1 genotypes were combined to determine the presence of DQ2.5, DQ2.2, DQ8, and DQ7.5 haplotypes (**Supplementary Table 1**). Frequencies of the imputed genotypes were in line with observed European CD patient and general population frequencies (**Supplementary Table 2**), further validating our approach[11]. Using imputed genotypes, samples were stratified into 15 genotypes representing all possible combinations of risk haplotypes. Counts for each HLA genotype in cases and controls for each population are detailed in **Supplementary Table 3**.

### Existing risk prediction models

HLA genotypes were grouped to match previously described approaches for HLA based risk stratification (**Supplementary Table 4**). The Romanos *(ROM)* [17] and Tye-Din *(TD)* [8] models were constructed by stratifying samples based on their HLA genotypes into 3 or 6 categories respectively, coded as integers either 1–3 or 1–6. The model developed by Abraham *et al. (GRS*_228_) is based on the weighted sum of 228 SNP calls, with the SNPs and weights derived from previous application of an L1-penalized support-vector machine (SVM) to the UK2 cohort[16].

### The *HDQ* model

We constructed two models that consider the possible risk HLA genotypes occurring at the HLA-DQA1/DQB1 locus: *HDQ_15_* based on the 15 possible genotypes of known risk haplotypes (**Supplementary Tables 4**) and *HDQ_17_* which considers a further two risk genotypes discovered in this work.

Using *G* to denote the genotype of the HLA-DQA1/DQB1 locus, and *Y* to denote the binary phenotype, the *HDQ* models can be expressed as

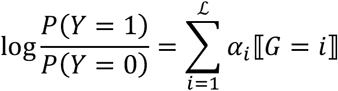

where 〚.〛 is the Iverson bracket, *G* = *i* indicates the *i*-th genotype at HLA-DQA1/DQB1 locus, with 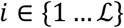 where 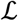 is 15 or 17 for *HDQ*_15_ and *HDQ*_17_ respectively (further details in the Supplementary Methods). Model weights were derived exclusively in the UK2 dataset. The ability of risk scores to generalize to unseen data could then be evaluated in the four other European populations.

### Identification of novel interactions with HLA risk alleles, conditioning on the HDQ_15_ model

To determine if further HLA risk factors could be identified beyond the *HDQ*_15_ model, we systematically evaluated *trans* interaction effects between HLA-DQA1 and HLA-DQB1 haplotypes in combination with the known risk haplotypes. The test considers the *HDQ*_15_ model and incorporates a novel dosage-encoded haplotype *X_j_*

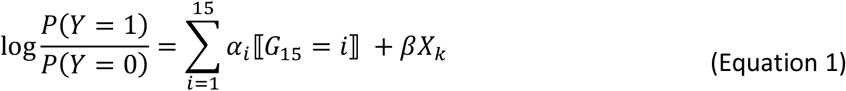

where *X_j_* ∈ {0,1,2}. This model is expanded to also including an interaction term

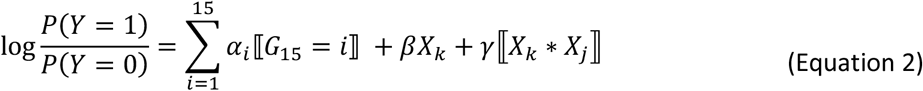

where *X_k_* is a dosage-encoded risk haplotype. We used a likelihood ratio test to compare the fit of models given by Equation 1 and 2 to assess the statistical significance of the interaction, with the test statistic following a *χ*^2^ distribution with 1 degrees of freedom. We only consider interactions where the common haplotype and its interaction with a known risk haplotype both have an allele frequency of at least 1% in the discovery (UK2) population, leading to 15 independent tests in total. Further details in **Supplementary Methods**.

### Identification of HLA haplotype SNP tags

SNP tags [20] were identified for all relevant haplotypes using the T1DGC and 1000 Genomes EUR populations (**Supplementary Methods**). The best performing available tags are detailed in **Supplementary Table 8-8**.

### Measures of Predictive Performance

AUC values were used as the primary measure to quantify predictive performance of all models. Significance of the AUC differences was evaluated using DeLong’s test for paired ROC curves (calculated using the R package pROC [22]).

### Data Visualisation

All plots were derived using R (version 3.3.2)[23] using the package ggplot2[24].

## Results

### Coeliac disease risk estimates from known risk haplotypes can achieve predictive power greater than previously indicated

The *ROM* model [17] has previously been used as the HLA-attributed risk prediction baseline against which novel GRS models are compared [16, 17]. Examining the distribution of risk across each of the 15 HLA-DQ genotypes (**Figure 1**) showed that while the *ROM* ‘High’ and ‘Low’ categories represent two clear extremes of CD risk, the risk attributed by ‘Intermediate’ categories were highly variable with ORs ranging from that of DQ2.2/DQ7.5 *(OR* = 1.35–5.89) to DQ2.2/DQX *(OR* = 0.09–0.28).

**Fig 1.**
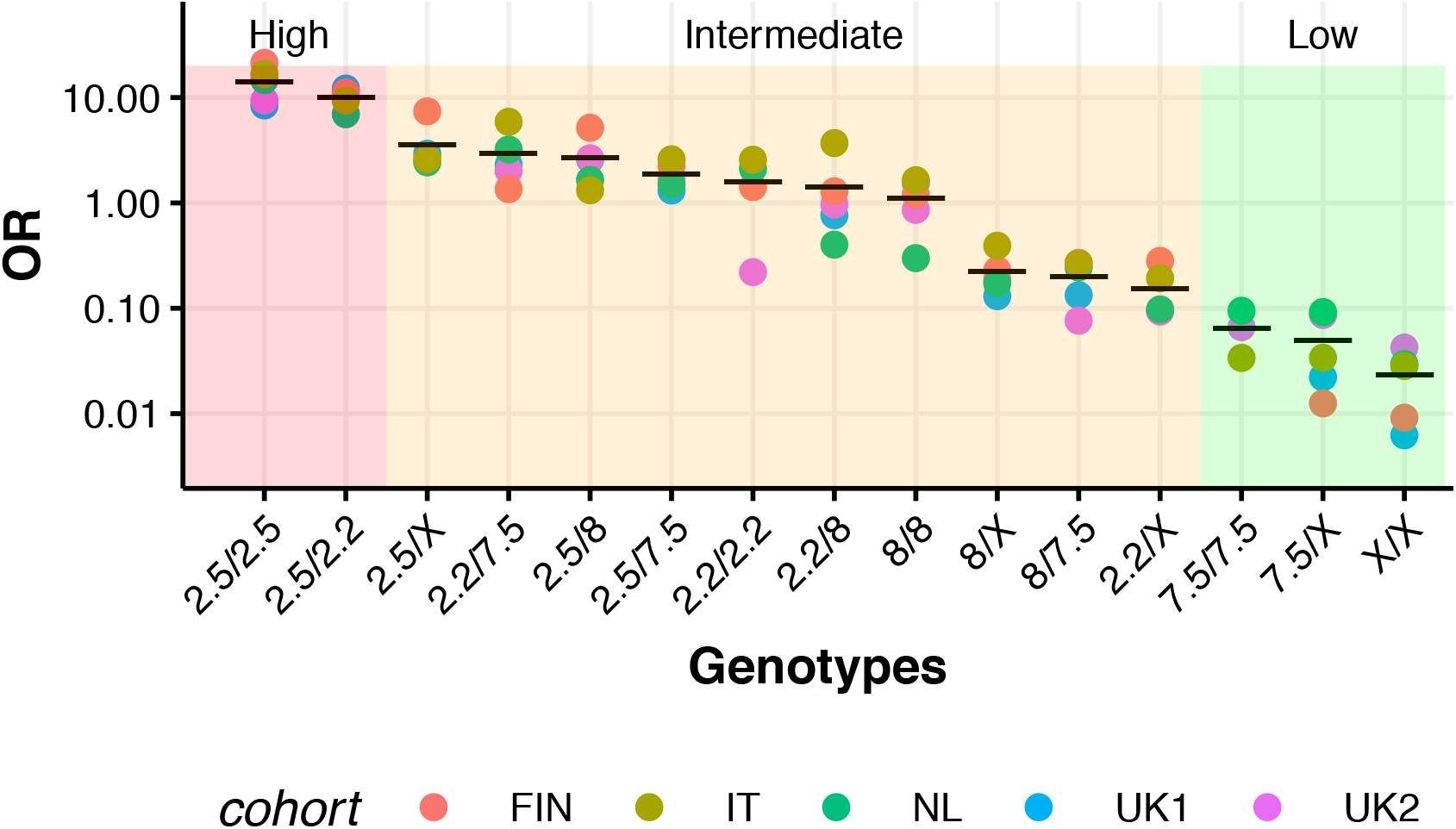
Effect sizes each of the known CD associated HLA-DQ risk genotypes. Predictive value as characterised by odds ratios (OR) for each of the 15 HLA-DQ risk genotype combinations in five European CD case/control populations. Genotypes are sorted by average OR of each genotype (horizontal bar). A OR of 0 was observed for DQ2.2/DQ2.2 and in UK1 and DQ7.5/DQ7.5 in FIN and UK1 as they were not observed in any CD carriers and hence are shown on this graph as “marginal” outlier dots. Red, orange and green shaded areas indicate which genotypes fall into the “high”, “intermediate” and “low” risk categories from risk model ROM by Romanos *et al.* [17].

As these results suggest that stratification of patients using the *ROM* model may not accurately represent HLA mediated risk, we constructed a risk score, *HDQ_15_,* using data from the UK2 population. Predictive performance of this model was assessed in 4 independent validation populations and a combined validation population alongside predictive performance of two previously reported HLA models, *ROM* and GT[8], and a state-of-the-art CD genomic risk score reported by Abraham *et al., GRS_228_* (**Figure 2A, Supplementary Table 10, Supplementary Figure 1**).

**Fig 2.**
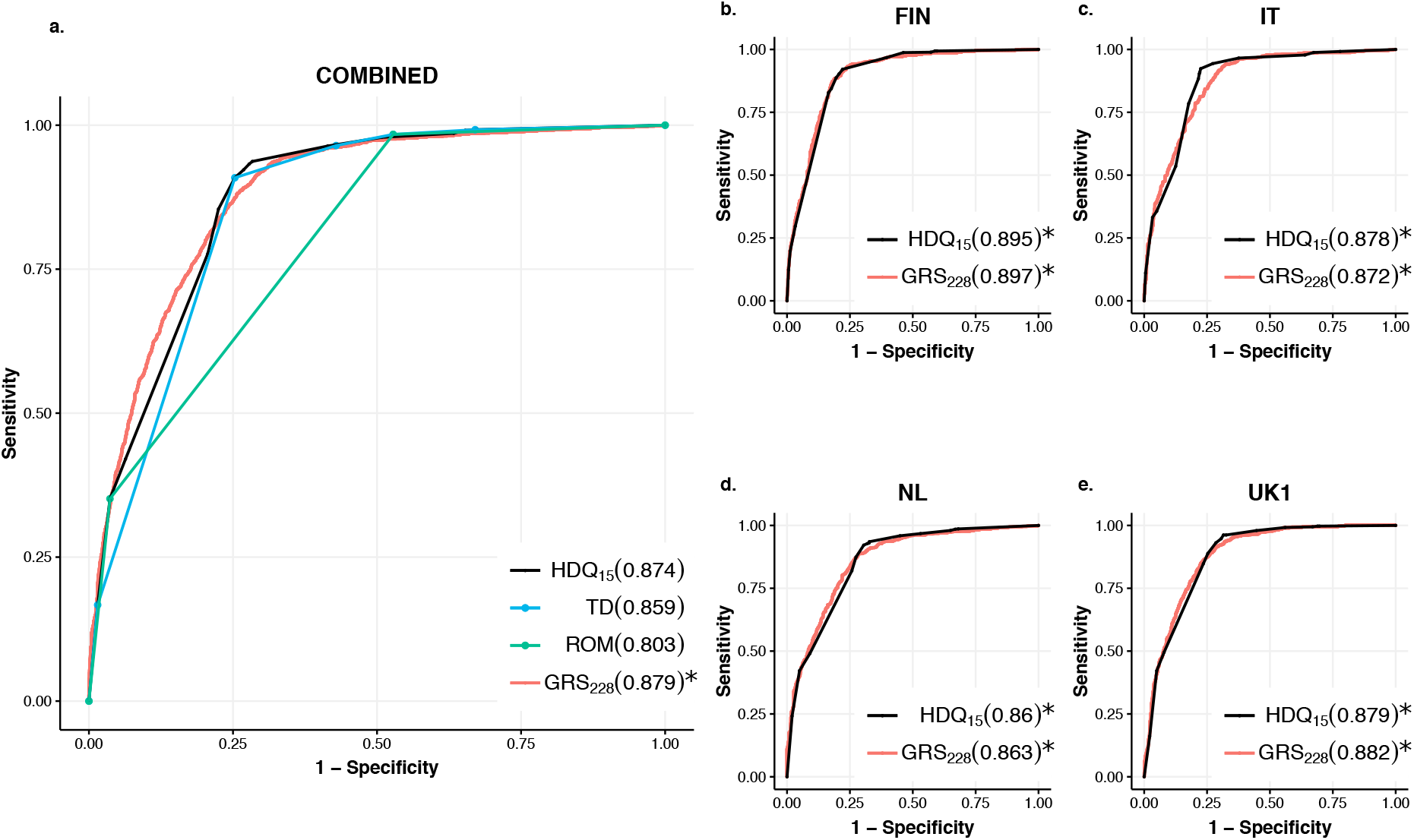
ROC curves of the CD risk models considered in this work in 4 external validation cohorts, combined or individually. Curves in **a)** represent the performance of the HDQ_15_, TD, ROM and *GRS*_228_ models over all validation cohorts combined. Curves on the right represent performance of HDQ_15_and GRS_228_ in the **b)** FIN, **c)** IT, **d)** NL and e) UK1 cohorts, with models trained using the UK2 dataset. The AUCs for each model is shown in the legend of each plot, and models that had the highest AUC or were not significantly different are marked with an asterisk.

In all populations, the *HDQ_15_* and *GRS_228_* models performed equivalently and significantly outperformed all other models. However, on the combination of all four test cohort, the increased performance of *GRS_228_* was significant *(AUC: 0.874 vs* 0.880 for *HDQ_15_ and GRS_228_ p = 0.0077).* The performance of the *ROM* model was much lower (~5%) than all other models in each of the validation populations. The *TD* model, recently recommended for clinical practice, performed only marginally worse than the best performing models (~2%). While the overall AUCs of the SNP-based *GRS_228_* and HLA haplotype based *HDQ_15_* were almost identical, differences could be observed between the shapes of ROC curves (**Figure 2B-E**).

### The DQ6.2 and DQ7.3 haplotypes modulate DQ2.5 risk and can improve CD prediction

Given the increasing evidence that interactions between HLA alleles can improve risk stratification in autoimmune diseases[14, 15], we sought to determine whether interactions between known and novel HLA-DQA1 – HLA-DQB1 risk haplotypes could be identified which further improve our predictive model.

An analysis of the interaction effects of 15 HLA-DQA1-HLA-DQB1 haplotypes with the known CD risk alleles conditioning on the 15 terms in the *HDQ_15_* model (**Supplementary Table 11**) found that two HLA-DQ haplotypes DQ6.2 (HLA-DQA1*01:02-DQB1*06:02) and DQ7.3 (HLA-DQA1*03:03-DQB1*03:01) show significant epistatic interactions with DQ2.5 (**Table 2**). DQ6.2 was found to increase risk when observed with DQ2.5 (OR=4.22 in UK2, OR=2.55 in validation) and DQ7.3 was (**Supplementary Table 10, Supplementary Figure 2**). found to decrease risk when observed with the DQ2.5 haplotype (OR=0.13 in UK2, OR=-0.47 in validation). These modulating effects could also be observed in each of the 4 validation populations (**Figure 3A**). Interestingly, DQ7.3 also showed significant additive in both the UK2 discovery and combined validation cohorts (**Table 2**).

**Table 2:** P-values and odds ratios for novel risk haplotypes DQ6.2 (HLA-DQA1*01:02-DQB1*06:02) and DQ7.3 (HLA-DQA1*03:03-DQB1*03:01) as additive effects (top half) or interacting with DQ2.5 (bottom half). Each analysis is further separated into the discovery phase from the UK2 dataset, and the replication results from all remaining cohorts combined. –log10(P) indicates the **-log_10_(P)** and OR is the odds ratio (and confidence interval).

|  |  | Discovery (UK2) |  | Replication(Comb.) |  |
| --- | --- | --- | --- | --- | --- |
| | Genotype | $-\log_{10}(P)$ | OR | $-\log_{10}(P)$ | OR |
| Additive | DQ6.2 | 9.97 | 1.94 (1.60-2.34) | 0.95 | 1.21 (1.01-1.45) |
|  | DQ7.3 | 9.97 | 0.32 (0.23-0.47) | 11.05 | 0.25 (0.17-0.39) |
| Interaction with DQ2.5 | DQ6.2/DQ2.5 | 8.63 | 4.22(2.61-6.82) | 3.48 | 2.55 (1.57-4.16) |
|  | DQ7.3/DQ2.5 | 6.71 | 0.13 (0.07-0.27) | 3.37 | 0.19 (0.09-0.42) |

When the DQ2.5/DQ6.2 and DQ2.5/DQ7.3 risk genotypes were added to the *HDQ*_15_ model to construct a 17-category model *(HDQ*_17_), predictive performance increased significantly in the combined validation population (AUC: *0.874* and 0.883 for *HDQ_15_* and *HDQ_17_*, p=7.84×10--12) (Supplementary Table 10). No significant difference in risk prediction could be observed between the *HDQ*_17_ and *GRS*_228_ on the combined validation (Figure 3B). To ensure that the results observed in this work were not driven by imputation error, we identified a set of six SNPs that could be used to establish the presence of HLA risk haplotypes and used these to re-implement the *HDQ*_17_ model. The predictive performance of the *HDQ_17_* model derived from SNP calls was not statistically different to the haplotypes determined from imputed HLA genotype, in all populations

**Fig 3:**
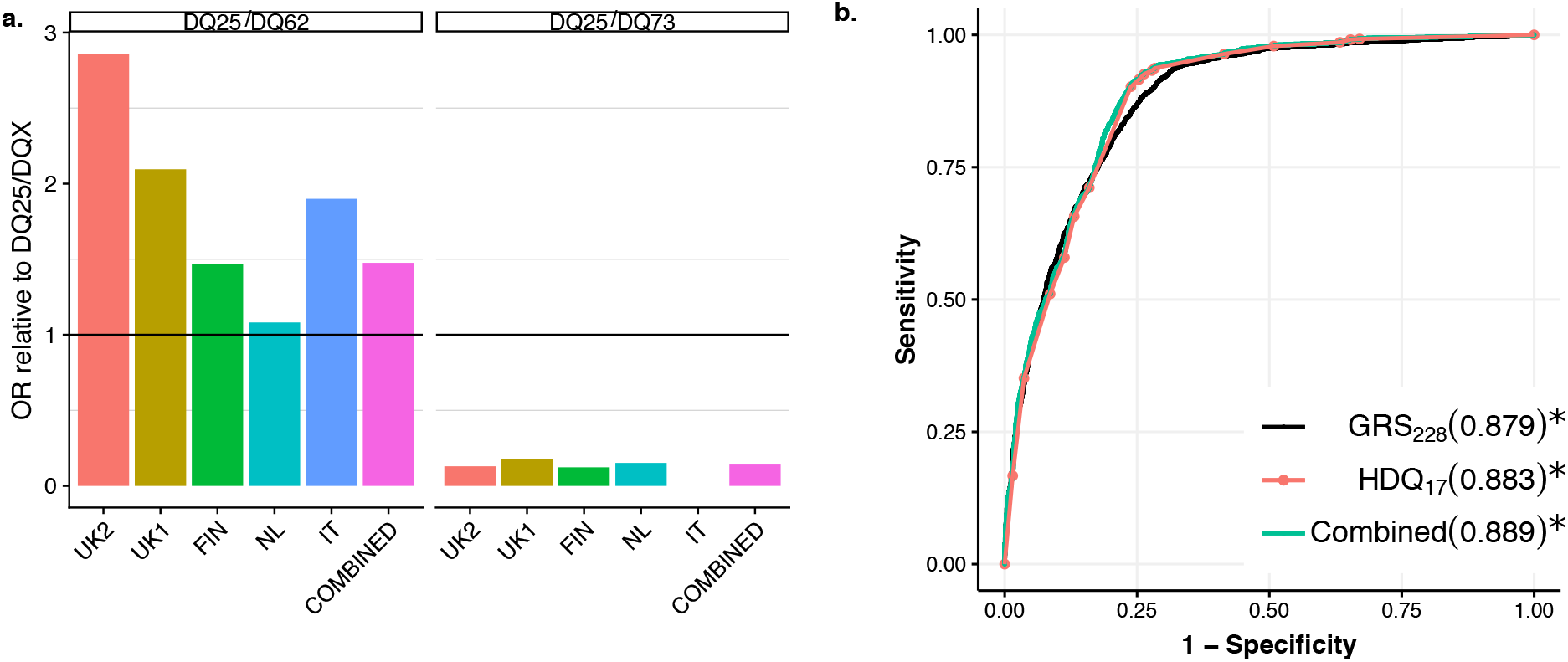
Effect size and impact on prediction of the novel interactions of DQ7.3 and DQ6.2. **a)** Effect size (OR) of the DQ6.2/DQ2.5 and DQ7.3/DQ2.5 relative to the DQ2.5/DQX genotype, showing the consistent deleterious and protective effects of these genotypes. The DQ7.3/DQ2.5 is not present in cases in the IT cohort. b) ROC curves for the over the combined validation cohorts, showing **b)** HDQ_17_ and GRS_228_ indicating the improvement in prediction contributed by the two novel interactions. **c)** the strong performance of HDQ_17_ that remains when implemented using tag SNPs and **d)** the small improvement that can be obtained when the HDQ_17_ and GRS_228_ are combined compared the individual models. Models that had the highest AUC or were not significantly different are marked with an asterisk.

### CD screening exclusionary criteria may be modified to improve predictive value using the *HDQ_17_* model

The improved risk stratification of the *HDQ_17_* model offers an opportunity to re-examine the existing exclusionary criteria used in CD genetic testing to determine whether the high negative predictive value (NPV) can be maintained while improving positive predictive values (PPV). As the three lowest risk haplotypes in the *HDQ*_17_ model correspond to the existing exclusionary criteria, we consider the impact of changing the exclusion criteria to also exclude individuals who fell into the 4^th^-6^th^ lowest risk *HDQ_17_* categories.

The impact of altering the exclusion criteria on the trade-off between PPV and NPV at a CD prevalence of 1% (approximate frequency in the general population) and 10% (approximate frequency in first degree relatives) is shown in **Figure 4**. At a population frequency of 1% (**Figure 4A**), a mean NPV above 0.999 can be maintained even if the 6 lowest HLA risk genotypes per the *HDQ_17_* model are treated as negative for CD. In contrast, PPV increased across all cohorts from a mean PPV of 1.9% for the “No DQ2/DQ8” exclusionary criteria to 3.3% for the 6 categories with the lowest CD-risk according to *HDQ_17_.* A similar effect is observed at a CD prevalence of 10% (**Figure 4B**), where the usage of these 6 categories as an exclusion cut-off yields PPV improvement from 17.5% to 27.1% while maintaining NPV above 99%. Such an increase in PPV has potential to reduce the number of patients requiring diagnostic follow-up when screening at-risk groups.

**Fig 4:**
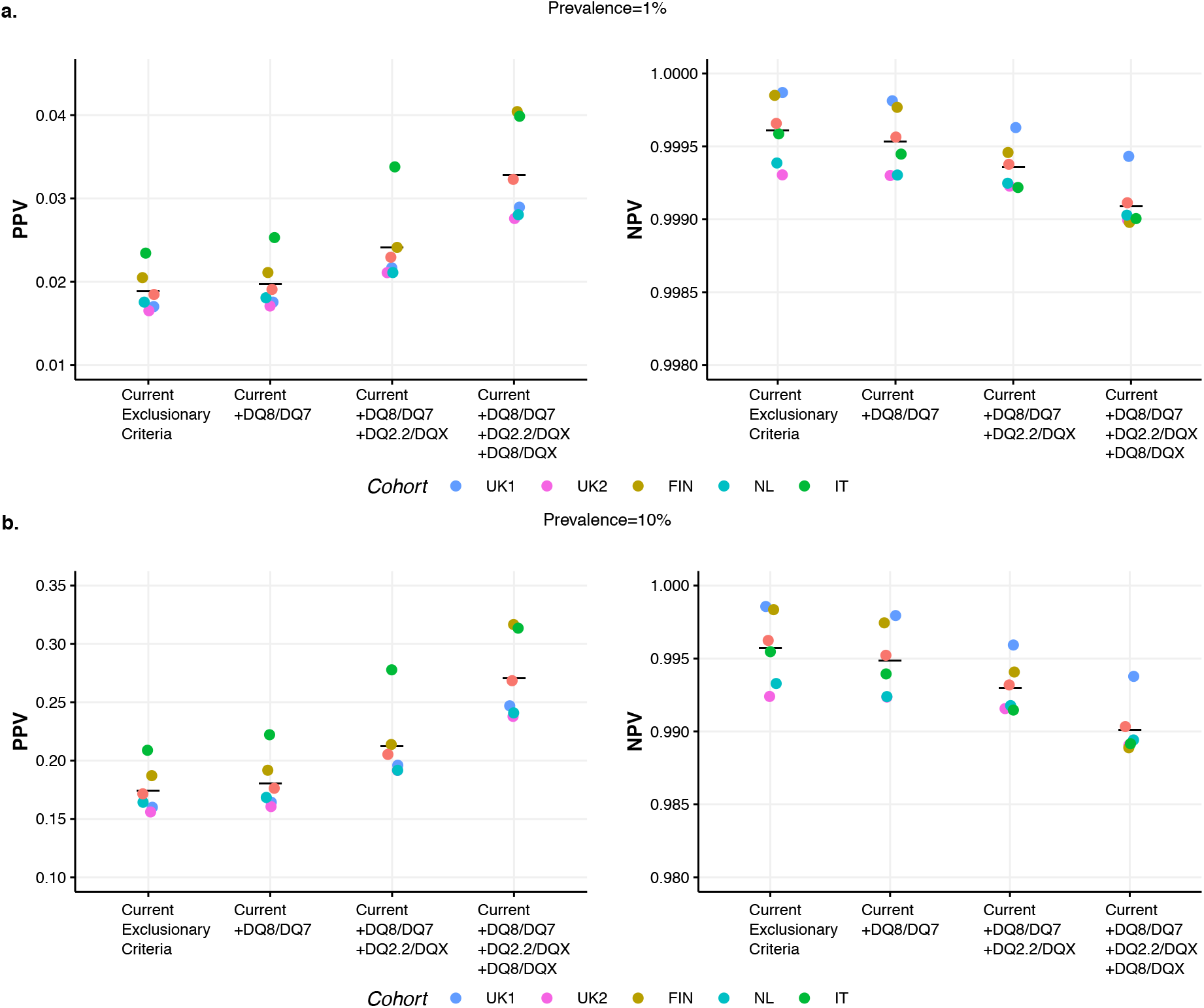
PPV and NPV of different CD exclusion criteria using the HDQ_17_ model. Boxplots show the impact of altering the diagnostic exclusionary criteria for CD diagnosis from the current diagnostic criteria (e.g. carrying no DQ2 or DQ8 risk alleles – DQX/DQX, DQX/DQ7.5, DQ7.5/DQ7.5) to also excluding DQ8/DQ7.5, DQ2.2/DQX and/or DQ8/DQX at a disease frequency of (A) 1% and (B) 10% (note the difference in scale on the y-axes). These additional genotypes were selected as they are based on the lowest categories of the HDQ_17_ model. Individual points within each category represents a different cohort and horizontal lines represent the mean.

## Discussion

Despite the high positive predictive value of serological testing and the high negative predictive value of HLA genotyping, biopsy via upper gastrointestinal endoscopy, a resource intensive, invasive and inconvenient process, remains the gold standard for diagnosis of CD. There is a need for improved risk stratification strategies to reduce unnecessary endoscopies and improve the efficiency of CD investigation, particularly in the screening of high-risk, asymptomatic individuals. This work demonstrates that the application of genomic risk score frameworks using only HLA risk haplotypes leads to a predictive performance that significantly outperforms current HLA stratification approaches and can be used to help further uncover the genetic basis of this condition.

A potential clinical impact of this research is the modification of the exclusionary criteria for CD when using genetic testing as a screening tool, especially for children at high risk for CD due to family history, but who are currently asymptomatic. For this at-risk group, recent European guidelines recommend an HLA testing as an initial screening test[26]. If we tested 100,000 individuals from this high-risk group and incorporated DQ8/DQ7.5, DQ2.2/DQX and DQ8/X genotypes into the exclusionary criteria, we would correctly exclude 64,279 individuals from a CD diagnosis and hence from biopsy, a further 21,249 individuals compared to the 43,030 using the current threshold (**Supplementary Table 12**). Similar increases in the number of true-negative diagnoses are observed with a population prevealance of 1% (70,706 up from 47,333) (**Supplementary Table 12**). While this shift in exclusionary criteria leads to a small increase in the false positive rate, this may be allieviated further monitoring or serological testing. As with varying the inclusion criteria for serology testing [26], further studies need to be conducted to better quantify the impact of any changes.

By conditioning on the *HDQ_15_* model, two novel haplotypes which modulate HLA risk may be identified which significantly improve HLA mediated prediction, and taken with previous studies provide further evidence that non-additive interactions between HLA loci may be common. While additional independent SNPs within the HLA region have been previously identified by CD GWAS[27], no study has translated this signal to associations with DQ6.2 or DQ7.3. The effect size of these variants in their interaction with DQ2.5 in the validation cohort are more extreme than the strongest reported non-HLA variants. Their impact on risk prediction is significant and indicates their potential importance for understanding the underpinnings of CD. We note however that there is clear variability across the different populations, especially for DQ6.2, indicating the potential for other modifiers of this relationship.

Interestingly, we found that DQ7.3 was significant in discovery and replication cohorts when analysed univariately. We believe this haplotype may have been missed in previous studies due the typing resolution and sample size needed to detect the effects of this relatively rare haplotype. Future study of the mechanism by which these interactions modulate DQ2.5 risk will help inform how HLA alleles mediate disease predisposition on the molecular level.

The ability to implement the *HDQ_17_* model using either HLA genotype or SNP-tags indicates that it may be a more easily translatable alternative to existing GRS approaches given that HLA genotyping is already in use in routine CD diagnosis. Exclusionary typing for CD is now one of the most common genetic tests performed each year in Australia[8], and if this typing has already been performed, the *HDQ_17_* model may be implemented at no additional expense, to better understand patient risk. An interesting area of relevant research is point-of-care SNP genotyping [28, 29], where the small panel of SNPs, combined with point-of-care serological tools[30], may provide a pathway towards immediate, confident CD exclusion at a low cost in the clinical setting.

The HLA-only *HDQ_17_* model performs equivalently to state-of-the-art genomic prediction models using both HLA and non-HLA information in all validation cohorts. This has several implications for risk stratification in CD and other auto-immune diseases. Firstly, the contribution of non-HLA variation for CD risk prediction is unclear, given that previous comparisons were against a baseline that did not make full use of HLA risk variation. [17]Secondly, SNP-based implementation of the *HDQ_17_* model is far more parsimonious than models requiring hundreds of SNPs and may be a demonstration of the inability existing regularised machine learning techniques to find the smallest subset of features that yield the best predictive performance, a known issues for some regularized models[31]. Finally, these results may also indicate that HLA genotypes may have been underutilised in construction of GRS for other auto-immune conditions and that further exploration of fine grained HLA data may yield improved risk stratification for other conditions.

## Conclusion

This analysis demonstrates that through systematic interpretation of HLA attributed risk, it is possible to improve risk prediction of CD using known and novel HLA risk haplotypes. The proposed *HDQ_17_* risk haplotype model performs equivalently to a state-of-the-art genomic risk model across multiple distinct European patient cohorts, but only uses information that is routinely collected in a clinical setting. The improved understanding may be useful for refining the HLA typing exclusionary criteria, especially in screening of children at high risk of CD. These results may allow for a more refined clinical pathway for screening and diagnosis of CD and act as the foundation for the development of improved genomic prediction models.

## Supporting Information Legends Supplementary Tables

**Supplementary Table 1. Haplotype Calling from HIBAG imputed HLA-DQA1 and HLA-DQB1 Alleles**

**Supplementary Table 2. HIBAG Imputed genotype frequency vs observed frequency** Observed case and control frequencies from Koskinen *et al.* 2009[11].

**Supplementary Table 3. Imputed genotype frequency in cases and controls for each population**

**Supplementary Table 4**. The Tye-Din (TD) and Romanos (ROM) and proposed *HDQ_15_* HLA based CD risk models examined in this study. HDQ_1S_ and HDQ_o_ are HLA stratification models proposed in this paper while Tye-Din (TD) and Romanos (ROM) describe existing HLA based risk stratification models. The risk scores for HDQ_1S_ were derived in the UK2 dataset. Rows are sorted based on risk in the TD model, then by HDQ_1S_. Numbers in brackets represent the rankings of categories in each model from highest to lowest risk.

**Supplementary Table 5**. Proposed *HDQ_17_* HLA based CD risk models examined in this study. The risk scores for HDQ_17_ were derived in the UK2 dataset.

**Supplementary Table 6**. Odds ratios *(OR)* for 15 HLA Genotypes for each of five populations considered in this paper. HLA genotypes are sorted as per the *HDQ_15_* model using the UK2 OR.

**Supplementary Table 7**: AUCs for the HDQ15 when weights are positive likelihood ratios (LR+, as in the main paper), where the weights are based on the odds ratios (OR) of each risk allele or where logistic regression (Log. Reg.) fitted to all risk alleles together. The results are identical using the Log Reg and LR+ models.

**Supplementary Table 8**. Performance of HLA haplotype SNP tags in the T1DGC and 1000 Genomes (EUR) datasets.

**Supplementary Table 9**: Tagging SNP to HLA haplotype conversion

**Supplementary Table 10**: AUCs and significance for all models discussed in the paper across all cohorts. In all cohorts except UK2 (used for training the model), we’ve indicated which models are not significantly different from the best (the combined HDQ17+GRS228 model). This varies a lot between cohorts both due to model performance as well as the samples size of the different studies.

**Supplementary Table 11**: Frequency, P-values and odds ratios for interactions between common (AF>1%) haplotypes and known CD risk factors for discovery (UK2) and validation (Combined) cohorts. Interactions were only included if the two haplotypes occurred in more than 1% of the UK2 population (Freq). OR indicates the odds ratio with lo and high indicating the associated confidence interval. P indicates the -log10(P-value) for the interaction based on a likelihood ratio test. Grey shading is used to separate groups of interactions with the same novel haplotype. Bolding is used to indicate the two interactions that are significant in the discovery and replication cohorts. The threshold for significance here is 0.05/15=0.0033

**Supplementary Table 12**: Impact of shifting exclusionary criteria if we consider 100,000 tests with prevalence of either 1% or 10%. The numbers below are based on the average positive and negative predictive values derived from each of the five cohorts in our study, as shown in Figure 4 in the main text. The exclusionary criteria (i.e. genotypes that indicate a negative CD diagnosis) are as follows: current is DQX/DQX DQ7/DQX, DQ7/DQ7, current+1 adds DQ8/DQ7, current+2 adds DQ2.2/DQX, current+3 adds DQ8/DQX, as per Figure 4 in the main document.

**Supplementary Figure 1**: ROC curves of all models considered in this paper across the FIN, IT, NL and UK1 cohorts. Each corresponds to a different model and the corresponding AUC is shown in the legend. Staring of the legend entry indicates that the model is statistically equivalent to the best performing model.

**Supplementary Figure 2**: Performance of proposed HLA models using either imputed haplotypes or 4-6 tag SNPs. Left and right subplots represent the *HDQ_15_* and *HDQ_17_* models respectively, with the set of points indicating AUCs if the model were based on imputed haplotypes (Hap.) or tag SNPs (SNP), 4 SNPs for *HDQ_15_* and 6 for *HDQ_17_.* No significant differences were found between the use of imputed haplotypes or SNPs for either model as evaluated by DeLong’s test. Horizontal lines represent the mean AUC.

**Supplementary Figure 3** To assess whether predictive performance could be further improved through combination of the *HDQ_17_* and *GRS_228_* models, the performance of models variably weighting each predictor were assessed. No significant improvement could be observed over either model using this combination approach. The combination presented in the paper makes use of a 0.9 weighting of HDQ17, meaning the resulting combination reorders the samples within each of the 17 HDQ17 categories by the GRS228.

