## Supplementary Materials for "Improved HLA-based prediction of coeliac disease identifies two novel HLA risk modifiers, DQ6.2 and DQ7.3"

### Supplementary Methods

#### $HDQ_{15}$ – the base-line CD risk model

To formally describe the $HDQ_{15}$ model, we use $G_{15}$ to denote mapping of each sample, $x$, to one of the 15 mutually exclusive genotype classes $i \in \{1, 2, \dots, 15\}$ of the HLA-DQA1/DQB1 locus listed in Supplementary Table 1. We use $Y$ to denote the phenotype, 1 for celiac cases and 0 for controls. The $HDQ_{15}$ model can then expressed as the standard logistic regression solution for

$$\log \frac{P(Y=1)}{P(Y=0)} = \sum_{i=1}^{15} \alpha_i [\![G_{15} = i]\!] \qquad \text{(Equation 1)}$$

where $[\![P]\!]$ is the Iverson bracket (defined as 1 if the preposition inside the brackets is true and 0 otherwise), $G\ = i$ indicates the $i$-th genotype at HLA-DQA1/DQB1 locus, $i \in \{1 \dots 15\}$ and $\alpha_i$ is the coefficient for the $i$ -th genotype. Note that there is no "bias" term used above as it is redundant, since our 15 genotype classes are mutually exclusive. This mutation exclusion implies that the maximal likelihood solution to the above logistic regression model can be written explicitly in terms of log odds

$$\alpha_i = \text{logit}\left(\widehat{\text{P}}(\text{Y}=1|G_{15}=i)\right) := \log \frac{\widehat{\text{P}}(\text{Y}=1|G_{15}=i)}{\widehat{\text{P}}(\text{Y}=0|G_{15}=i)} = \log \frac{n_{1i}}{n_{0i}}. \qquad \text{(Equation 2)}$$

where $n_{1i}$ and $n_{0i}$ denote the numbers of cases and controls carrying the $i^{th}$ genotype, respectively.

Given this logistic regression model, we can express the risk of for a sample $x$ of the genotype class $i_x := G_{15}(x)$ as:

$$\text{p}_{15}(\text{x}) = \widehat{\text{P}}(Y = 1 \mid G_{15} = i_x) = \frac{1}{1 + \exp\left(-f_{15}(x)\right)} \qquad \text{(Equation 3)}$$

where the $f_{15}$ is the $HDQ_{15}$ logistic regression scoring function explicitly defined as

$$f_{15}\ (x) := \sum_{i=1}^{15} \alpha_i [\![G_{15} = i_x]\!] \ = \alpha_{G_{15}(x)} = \alpha_{i_x} \qquad \text{(Equation 4)}$$

With the coefficients $\alpha_i$ are given in the Supplementary Table 4.

The scoring function $f_{15}$ maximises the Area Under the ROC Curve (AUC) in the training (UK2) dataset[21] for all classifiers $f$ preserving all genotype classes $\{G_{15} = i\}$ . For such classifiers the AUC is dependent only on the ordering of those classes according to values allocated by $f$ and the maximal value is also achievable for the order of those classes according to the (empirical) positive likelihood ratios (PRL)

$$\lambda_i := \frac{\widehat{\mathrm{P}}(G_{15} = i|\mathrm{Y} = 1)}{\widehat{\mathrm{P}}(G_{15} = i|\mathrm{Y} = 0)} = \frac{n_{1i}n_{0+}}{n_{0i}n_{1+}}, \qquad \text{(Equation 4)}$$

for $i = 1,2,\dots,15$ and $n_{1+}$ and $n_{0+}$ denoting the total numbers of cases and controls in the data, respectively. The corresponding scoring model can be written as

$$f_{15}^{*}\ (x) = \textstyle\sum_{i=1}^{15} \lambda_i [\![G_{15} = i_x]\!] = \lambda_{i_x} \approx \frac{\widehat{\mathrm{P}}(G_{15} = i_x|\mathrm{Y} = 1)}{\widehat{\mathrm{P}}(G_{15} = i_x|\mathrm{Y} = 0)} \qquad \text{(Equation 5)}$$

for the sample $x$ of the genotype class $i_x := G_{15}(x)$. The coefficients $\lambda_i$ are given in Supplementary Table 4.

In this paper, all predictive models were derived exclusively using the UK2 dataset. The ability of risk scores to generalize to unseen data had been assessed by evaluation in the four other European populations.

### Identification of novel interactions with HLA risk alleles, conditioning on the $\mathrm{HDQ}_{15}$ model

Given the differences between the $HDQ_{15}$ and $GRS_{228}$ models, we hypothesised that there may be other HLA-DQA1/DQB1 haplotypes which could improve performance of our risk model. Given that additive effects have previously been explored, we have decided to evaluate systematically *trans* interaction effects for additional haplotypes.

From our imputed HLA data, we observed 37 haplotypes formed loci formed from HLA-DQA1 and HLA-DQB1 alleles, with many occurring at a very low frequency (<1%). As we are considering the interaction of novel haplotypes with known CD risk haplotypes , we only considered interactions such that the novel allele and the interaction between the novel allele and a known risk allele occur in $\geq 1\%$ samples in the UK2 cohort used for discovery. We have found 15 interactions, which satisfied this requirement, listed in Supplementary Table 10. These cover 6 potentially novel risk haplotypes and the 4 known risk haplotypes.

We refer to the 15 interactions to be analysed by the index, $G_{NE} \in \{1,2, \ldots ,15\}$. Alternatively, we can consider each of those genotypes is defined uniquely by a pair of dosage encoded haplotypes, $X$ where $X_j$, $j \in \{1,2, \ldots ,6\}$ refers to the six candidate haplotypes not previously associated with CD and $X_k$, $k \in \{1, \ldots 4\}$ refers to the four known risk haplotypes (see Supplementary Tables 1, 10 & 11 for details).

For each of those 15 novel genotypes, we have performed a separate interaction test using a likelihood ratio test. For the $l$-th genotype, with $G_{NE} = l \in \{1, \dots, 15\}$ or the corresponding pair of the $j$-th candidate and the $k$-th known risk haplotypes, we evaluate the fit of the baseline HDQ15 model (Eqn. 1) incorporating with the candidate dosage-encoded haplotype $X_j$

$$\log\frac{P(Y=1)}{P(Y=0)} = \sum_{i=1}^{15} \alpha_i [\![G_{15} = i]\!] + \beta X_k \quad \text{(Equation 6)}$$

where $X_j \in \{0,1,2\}$. This model is then compared with an expanded model including an interaction term

$$\begin{aligned}\log\frac{P(Y=1)}{P(Y=0)} &= \sum_{i=1}^{15} \alpha_i [\![G_{15} = i]\!] + \beta X_k + \gamma [\![G_{NE=j}]\!] \\ &= \sum_{i=1}^{15} \alpha_i [\![G_{15} = i]\!] + \beta X_j + \gamma [\![X_k * X_j]\!]\end{aligned} \quad \text{(Equation 7)}$$

where $X_k$ is a dosage-encoded known risk haplotype and $X_k * X_j$ indicates a given interaction of a novel and known risk haplotype. Both models (Eqns. 5 & 6) are solved in the standard logistic regression fashion by maximising maximal likelihood on the UK2 dataset. (Note we do not include here the redundant bias term; the values $X_k = 2$ may also be for some samples who have the DQX/DQX genotype).

A likelihood ratio test with 1 degree of freedom is applied to those solutions in order to establish whether the addition of an interaction term $\gamma [\![G_{NE=j}]\!] = \gamma [\![X_k * X_j]\!]$ resulted in a significant improvement of model's fit. In other words, we have performed a test of null hypothesis $\mathrm{H_0}: \gamma = 0$ versus alternative $\mathrm{H_1}: \gamma \neq 0$. The results in Supplementary Table 10 show that 2 out of total 15 trials showed a significant improvement, with DQ2.5/DQ6.2 & DQ2.5/DQ7.3 indicating a significant non-additive effect impact on CD risk .

#### $HDQ_{17}$ – an improved CD-risk model

Our next step is to develop new predictor by simply adding to the base-line $HDQ_{15}$ model, Eqn. 1, two additional terms representing two novel genotypes. The two risk interactions formed by DQ2.5/DQ6.2 & DQ2.5/DQ7.3 are incorporated as additional genotypes to the 15 that were already identified. Then, as in the case of $HDQ_{15}$, we can have the closed form solution to the logistic regression model:

$$\log\frac{P(Y=1)}{P(Y=0)} = \sum_{j=1}^{17} \beta_j [\![ G_{17} = j ]\!] \qquad \text{(Equation 8)}$$

As before the coefficients of the model fitted by the maximisation of likelihood can be written in the closed form:

$$\beta_j = \text{logit}\left(\widehat{\mathrm{P}}(\mathrm{Y}=1|G_{17}=j)\right) := \log\frac{\widehat{\mathrm{P}}(\mathrm{Y}=1|G_{17}=j)}{\widehat{\mathrm{P}}(\mathrm{Y}=0|G_{17}=j)} = \log\frac{n_{1j}}{n_{0j}} \qquad \text{(Equation 9)}$$

where $n_{1j}$ and $n_{0j}$ denote the number of cases and controls carrying the $j^{th}$ genotype in the dataset . The novel scoring function is

$$f_{17}(x) := \sum_{j=1}^{17} \beta_i [\![ G_{17} = j ]\!] = \beta_{G_{17}(x)} \qquad \text{(Equation 10)}$$

and the predicted probability of being the case

$$\mathrm{p}_{17}(x) = \widehat{\mathrm{P}}(Y=1 \mid G_{17}=j_x) = \frac{1}{1+\exp\left(-f_{17}(x)\right)} \qquad \text{(Equation 11)}$$

for any sample $x$ such that $G_{17}(x) = j_x$, see Supplementary Table 13 for more detail.

#### Existing risk prediction models

HLA genotypes were grouped to match previously described approaches for HLA based risk stratification (**Supplementary Table 5**). The Romanos ($ROM$) model was constructed by stratifying samples as High (DQ2.5/DQ2.5 or DQ2.5/DQ2.2), Low (DQ7.5/DQ7.5, DQ7.5/DQX or DQX/DQX) or Intermediate (all other categories) risk[17]. The Tye-Din ($TD$) model was constructed by stratifying samples into 6 categories; Highest (DQ2.5/DQ2.5), High (DQ2.5 positive), Moderate (DQ8 positive), Low (DQ2.2 positive), Very Low (DQ7.5 positive) and Lowest (DQX/DQX) based on relative-risk estimates in European populations [8]. These ordinal categories were coded as integers either 1–3 or 1–6 from highest to lowest risk for the $ROM$ and $TD$ models respectively.

The model developed by Abraham *et al.* ($GRS_{228}$) is the state-of-the-art for CD risk prediction. This model is based on the weighted sum of 228 SNP calls (in a dosage representation as values 0, 1 or 2, respectively), with the SNPs and weights derived from previous application of an L1-penalized support-vector machine (SVM) to the UK2 cohort[16]. As the details for this model are publicly available, we could determine the risk scores as detailed by Abraham *et al.* for each sample

#### Identification of HLA haplotype SNP tags

Inferring CD risk haplotypes using SNP tags is a well-established technique[11], however previously described SNP tags for the DQ2.2, DQ8 and DQ7.5 haplotypes were not available in the analysed datasets. To identify alternative SNP tags, an exhaustive comparison of all ~120,000 polygenic MHC SNPs in the 1000 Genomes dataset and each of the six risk haplotypes used in this work (DQ2.5, DQ2.2, DQ8, DQ7.5, DQ6.2 and DQ7.3) was performed in the 1,000 Genomes EUR population and T1DGC reference panel[22]. The best performing available tags are detailed in Supplementary Table 7. Samples were excluded from analysis if tag SNP genotypes were missing (0.5% of samples) or more than 2 HLA-DQ haplotypes tagged (0.1% of samples). A set of rules was manually derived to convert SNP genotype to HLA-DQ genotype (**Supplementary Table 8**).

### Supplementary Tables

**Supplementary Table 1. Haplotype Calling from HIBAG imputed HLA-DQA1 and HLA-DQB1 Alleles**

| HLA-DQA1 | HLA-DQB1 | HLA Haplotype |
|---|---|---|
| 05:01 | 02:01 | **DQ2.5** |
| 02:01 | 02:02 | **DQ2.2** |
| 03:01/02/03 | 03:02 | **DQ8** |
| 05:03/05:05 | Any | **DQ7** |
| All other | | **DQX** |

**Supplementary Table 2: HIBAG Imputed genotype frequency vs observed frequency.** Observed case and control frequencies from Koskinen *et al.* 2009.

| Sample Grouping | Imputed Case Freq. | Observed Case Freq. | Imputed Control Freq. | Observed control Freq. |
|---|---|---|---|---|
| DQ2.5 | 90.43% | 88.4-97.2% | 26.79% | 17.6-36.1% |
| DQ2.2 or DQ8 | 7.46% | 2.3-9.7% | 28.58% | 19.8-29.0% |
| DQ7 | 0.90% | 0-1.3% | 12.73% | 10.2-35.1% |
| DQX | 1.21% | 0-1.3% | 31.90% | 15.3-43.2% |

**Supplementary Table 3: Imputed genotype frequency in cases and controls for each population**

| | FIN | | IT | | NL | | UK1 | | UK2 | |
|---|---|---|---|---|---|---|---|---|---|---|
| **HLA Genotype** | **Cases** | **Controls** | **Cases** | **Controls** | **Cases** | **Controls** | **Cases** | **Controls** | **Cases** | **Controls** |
| DQ2.5/DQ2.5 | 80 | 11 | 55 | 3 | 194 | 17 | 119 | 57 | 314 | 104 |
| DQ2.5/DQ2.2 | 48 | 12 | 110 | 15 | 145 | 25 | 192 | 72 | 467 | 160 |
| DQ2.5/DQQ8 | 64 | 37 | 12 | 9 | 55 | 35 | 54 | 76 | 121 | 126 |
| DQ2.5/DQ7 | 28 | 35 | 49 | 21 | 38 | 26 | 31 | 83 | 81 | 129 |
| DQ2.5/DQX | 344 | 242 | 89 | 41 | 264 | 141 | 260 | 406 | 600 | 786 |
| DQ2.2/DQ2.2 | 1 | 1 | 7 | 2 | 4 | 1 | 0 | 28 | 5 | 59 |
| DQ2.2/DQ8 | 10 | 21 | 10 | 2 | 5 | 12 | 12 | 54 | 41 | 112 |
| DQ2.2/DQ7 | 10 | 20 | 124 | 28 | 44 | 14 | 30 | 46 | 77 | 103 |
| DQ2.2/DQX | 13 | 123 | 11 | 56 | 7 | 69 | 9 | 296 | 22 | 564 |
| DQ8/DQ8 | 11 | 25 | 3 | 1 | 2 | 6 | 11 | 24 | 20 | 61 |
| DQ8/DQ7 | 3 | 30 | 5 | 19 | 4 | 16 | 2 | 51 | 3 | 102 |
| DQ8/DQX | 30 | 319 | 10 | 26 | 19 | 103 | 13 | 313 | 45 | 576 |
| DQ7/DQ7 | 0 | 14 | 2 | 57 | 1 | 10 | 0 | 19 | 1 | 39 |
| DQ7/DQX | 1 | 200 | 6 | 143 | 10 | 101 | 2 | 283 | 18 | 503 |
| DQX/DQX | 4 | 739 | 4 | 120 | 11 | 270 | 2 | 788 | 34 | 1512 |
| **Total** | **647** | **1829** | **497** | **543** | **803** | **846** | **737** | **2596** | **1849** | **4936** |

**Supplementary Table 4. The Tye-Din (TD) and Romanos (ROM) and proposed $HDQ_{15}$ HLA based CD risk models examined in this study.** HLA based CD risk models examined in this study. $\mathrm{HDQ}_{15}$ and $\mathrm{HDQ}_4$ are HLA stratification models proposed in this paper while Tye-Din (TD) and Romanos (ROM) describe existing HLA based risk stratification models. The risk scores for $\mathrm{HDQ}_{15}$ were derived in the UK2 dataset. Rows are sorted based on risk in the TD model, then by $\mathrm{HDQ}_{15}$. Numbers in brackets represent the rankings of categories in each model from highest to lowest risk.

<table>
<tr><th rowspan="2">Genotype</th><th colspan="2">Existing</th><th>Proposed</th></tr>
<tr><th>$ROM$</th><th>$TD$</th><th>$HDQ_{15}$ score</th></tr>
<tr><td>DQ2.5/DQ2.5</td><td rowspan="2">High (1)</td><td>Highest (1)</td><td>1.105 (1)</td></tr>
<tr><td>DQ2.5/DQ2.2</td><td rowspan="5">High (2)</td><td>1.071 (2)</td></tr>
<tr><td>DQ2.5/DQ8</td><td rowspan="10">Intermediate (2)</td><td>-0.040 (3)</td></tr>
<tr><td>DQ2.5/DQX</td><td>-0.270 (4)</td></tr>
<tr><td>DQ2.2/DQ7</td><td>-0.291 (5)</td></tr>
<tr><td>DQ2.5/DQ7</td><td>-0.465 (6)</td></tr>
<tr><td>DQ8/DQ2.2</td><td rowspan="4">Moderate (3)</td><td>-1.005 (7)</td></tr>
<tr><td>DQ8/DQ8</td><td>-1.115 (8)</td></tr>
<tr><td>DQ8/DQX</td><td>-2.549 (10)</td></tr>
<tr><td>DQ8/DQ7</td><td>-3.526 (13)</td></tr>
<tr><td>DQ2.2/DQ2.2</td><td rowspan="2">Low (4)</td><td>-2.468 (9)</td></tr>
<tr><td>DQ2.2/DQX</td><td>-3.244 (11)</td></tr>
<tr><td>DQ7/DQX</td><td rowspan="3">Low (3)</td><td rowspan="2">Very Low (5)</td><td>-3.330 (12)</td></tr>
<tr><td>DQ7/DQ7</td><td>-3.664 (14)</td></tr>
<tr><td>DQX/DQX</td><td>Lowest (6)</td><td>-3.795 (15)</td></tr>
</table>

**Supplementary Table 5. Proposed $HDQ_{17}$ HLA based CD risk models examined in this study.** The risk scores for $\mathrm{HDQ}_{17}$ were derived in the UK2 dataset.

| Genotype | Proposed $HDQ_{17}$ score |
|---|---|
| **DQ2.5/DQ2.5** | 1.11 |
| **DQ2.5/DQ2.2** | 1.07 |
| **DQ25/DQ6.2** | 0.45 |
| **DQ2.5/DQ8** | -0.04 |
| **DQ2.5/DQX** | -0.29 |
| **DQ2.2/DQ7** | -0.44 |
| **DQ2.5/DQ7** | -0.47 |
| **DQ8/DQ2.2** | -1.00 |
| **DQ8/DQ8** | -1.12 |
| **DQ25/DQ7.3** | -2.16 |
| **DQ8/DQX** | -2.47 |
| **DQ8/DQ7** | -2.55 |
| **DQ2.2/DQ2.2** | -3.24 |
| **DQ2.2/DQX** | -3.33 |
| **DQ7/DQX** | -3.53 |
| **DQ7/DQ7** | -3.66 |
| **DQX/DQX** | -3.79 |

**Supplementary Table 6: Odds ratios ($OR$) for 15 HLA Genotypes for each of five populations considered in this paper.** HLA genotypes are sorted as per the $HDQ_{15}$ model using the UK2 **OR**.

| HLA Genotype | FIN | IT | NL | UK1 | UK2 | Average |
|---|---|---|---|---|---|---|
| DQ25/DQ22 | 11.18 | 9.36 | 6.95 | 12.16 | 10.02 | 9.93 |
| DQ25/DQ25 | 21.34 | 16.76 | 14.65 | 8.42 | 9.41 | 14.11 |
| DQ25/DQ8 | 5.17 | 1.32 | 1.65 | 2.58 | 2.65 | 2.68 |
| DQ25/DQX | 7.39 | 2.60 | 2.43 | 2.93 | 2.53 | 3.58 |
| DQ22/DQ7 | 1.35 | 5.89 | 3.21 | 2.30 | 2.02 | 2.95 |
| DQ25/DQ7 | 2.25 | 2.59 | 1.51 | 1.31 | 1.69 | 1.87 |
| DQ22/DQ8 | 1.29 | 3.70 | 0.40 | 0.76 | 0.97 | 1.42 |
| DQ8/DQ8 | 1.20 | 1.64 | 0.30 | 1.56 | 0.86 | 1.11 |
| DQ22/DQ22 | 1.41 | 2.57 | 2.11 | NA | 0.22 | 1.58 |
| DQ8/DQX | 0.23 | 0.39 | 0.17 | 0.13 | 0.19 | 0.22 |
| DQ22/DQX | 0.28 | 0.19 | 0.10 | 0.10 | 0.09 | 0.15 |
| DQ7/DQX | 0.01 | 0.03 | 0.09 | 0.02 | 0.09 | 0.05 |
| DQ8/DQ7 | 0.27 | 0.27 | 0.24 | 0.13 | 0.08 | 0.20 |
| DQ7/DQ7 | NA | 0.03 | 0.09 | NA | 0.07 | 0.06 |
| DQX/DQX | 0.01 | 0.03 | 0.03 | 0.01 | 0.04 | 0.02 |

**Supplementary Table 7:** AUCs for the $HDQ_{15}$ when weights are positive likelihood ratios (LR+, as in the main paper), where the weights are based on the odds ratios (OR) of each risk allele or where logistic regression (Log. Reg.) fitted to all risk alleles together. The results are identical using the Log Reg and LR+ models.

| | **FIN** | **IT** | **NL** | **UK1** | **UK2** | **Combined** |
|---|---|---|---|---|---|---|
| OR | 0.8946 | 0.8761 | 0.8565 | 0.8805 | 0.8611 | 0.8735 |
| Log. Reg. | 0.8950 | 0.8779 | 0.8600 | 0.8793 | 0.8613 | 0.8742 |
| LR+ | 0.8950 | 0.8779 | 0.8600 | 0.8793 | 0.8613 | 0.8742 |

**Supplementary Table 8. Performance of HLA haplotype SNP tags in the T1DGC and 1000 Genomes (EUR) datasets**

| | | T1DGC | | | | 1000 Genomes (EUR) | | | |
|---|---|---|---|---|---|---|---|---|---|
| **Haplotype/Allele** | **SNP Tag** | **D’** | **$R^2$** | **Sensitivity** | **Specificity** | **D’** | **$R^2$** | **Sensitivity** | **Specificity** |
| DQ2.5* | rs2187668_T | 0.997 | 0.975 | 99.8% | 99.5% | 1.000 | 0.982 | 100.0% | 99.8% |
| DQ2.2^ | rs2856705_A | 0.992 | 0.753 | 99.3% | 98.0% | 1.000 | 0.903 | 100.0% | 99.1% |
| DQ8[$] | rs9275312_G | SNP Not Available in T1DGC dataset | | | | 1.000 | 0.536 | 100.0% | 91.1% |
| DQA1*05:XX | rs3129763_T | 0.986 | 0.956 | 97.9% | 99.6% | Allele Not Available in 1000 Genomes dataset | | | |
| DQ6.2[#] | rs9271366_C | 0.941 | 0.586 | 94.6% | 97.2% | 0.986 | 0.862 | 98.7% | 98.2% |
| DQ7.3[&] | rs9357152_C | 0.952 | 0.551 | 95.7% | 92.9% | 0.987 | 0.634 | 99.0% | 91.7% |

*DQA1 alleles not annotated in 1000 Genomes dataset. Instead the DRB1*03:01-DQB1*02:01 haplotype is used to represent DQ2.5

^DQA1 alleles not annotated in 1000 Genomes dataset. Instead the DRB1*07:01-DQB1*02:01 haplotype is used to represent DQ2.2

[$] DQA1 alleles not annotated in 1000 Genomes dataset. Instead the DQB1*03:02 allele is used to represent DQ8

[#]DQA1 alleles not annotated in 1000 Genomes dataset. Instead the DQB1*06:02 allele is used to represent DQ6.2

[&]DQA1 alleles not annotated in 1000 Genomes dataset. Instead the DQB1*03:01 allele is used to represent DQ7.

**Supplementary Table 9: Tagging SNP to HLA haplotype conversion matrix**

| DQA1*05 | DQ2.5 | DQ2.2 | DQ8 | DQB1*03:01 | DQB1*06:02 | Genotype |
|---|---|---|---|---|---|---|
| rs3129763_T | rs2187668_T | rs2856705_A | rs9275312_G | rs9357152_C | rs9271366_C | |
| ≤2 | 2 | 0 | 0 | - | - | DQ2.5/DQ2.5 |
| ≤1 | 1 | 1 | 0 | - | - | DQ2.5/DQ2.2 |
| ≤1 | 1 | 0 | 1 | - | - | DQ2.5/DQ8 |
| 2 | 1 | 0 | 0 | - | - | DQ2.5/DQ7.5 |
| ≤1 | 1 | 0 | 0 | 0 | 1 | DQ2.5/DQ6.2 |
| ≤1 | 1 | 0 | 0 | 0 | 0 | DQ2.5/DQX |
| ≤1 | 1 | 0 | 0 | 1 | 0 | DQ2.5/DQ7.3 |
| 0 | 0 | 2 | 0 | - | - | DQ2.2/DQ2.2 |
| 0 | 0 | 1 | 1 | - | - | DQ2.2/DQ8 |
| 1 | 0 | 1 | 0 | - | - | DQ2.2/DQ7.5 |
| 0 | 0 | 1 | 0 | - | - | DQ2.2/DQX |
| 0 | 0 | 0 | 2 | - | - | DQ8/DQ8 |
| 1 | 0 | 0 | 1 | - | - | DQ8/DQ7.5 |
| 0 | 0 | 0 | 1 | - | - | DQ8/DQX |
| 2 | 0 | 0 | 0 | - | - | DQ7.5/DQ7.5 |
| 1 | 0 | 0 | 0 | - | - | DQ7.5/DQX |
| 0 | 0 | 0 | 0 | - | - | DQX/DQX |

**Supplementary Table 10:** AUCs for all models discussed in the paper across all cohorts and p-values indicating how different AUCs are from a given model. AUCs for each model is shown in a grey row, with significance of AUC different shown in the unshaded rows below. In all cohorts except UK2 (used for training the model), we've indicated (bolded) which AUCs are not significantly different from the best (the combined $HDQ_{17}$+$GRS_{228}$ model). This varies between cohorts due to both model performance and the samples size of the different studies. The significance tests for each model one-sided Delong test p-values for models against the combined $HDQ_{17}$ + $GRS_{228}$, $HDQ_{17}$ and $GRS_{228}$. (Bonferroni corrected significance threshold for the tests are 0.05/5, 0.05/4 and 0.05/3 respectively). Each model was only tested against those models which showed improved AUCs.

| | Combined | FIN | IT | NL | UK1 | UK2 |
|---|---|---|---|---|---|---|
| $HDQ_{17}$+ $GRS_{228}$ | **0.889** | **0.900** | **0.892** | **0.870** | **0.895** | 0.888 |
| $HDQ_{17}$ | **0.883** | **0.897** | **0.890** | **0.866** | **0.889** | 0.873 |
| *P-value vs $HDQ_{17}$ + $GRS_{228}$* | **0.11** | **0.32** | **0.4** | **0.36** | **0.22** | - |
| $GRS_{228}$ | **0.880** | **0.897** | **0.872** | **0.863** | **0.882** | 0.901 |
| *P-value vs $HDQ_{17}$ + $GRS_{228}$* | **0.023** | **0.32** | **0.063** | **0.27** | **0.044** | - |
| *P-value vs $HDQ_{17}$* | **0.07** | **0.49** | **0.016** | **0.28** | **0.043** | - |
| $HDQ_{15}$ | 0.874 | **0.895** | **0.878** | **0.860** | **0.879** | 0.861 |
| *P-value vs $HDQ_{17}$ + $GRS_{228}$* | 0.00073 | **0.25** | **0.15** | **0.17** | **0.016** | - |
| *P-value vs $HDQ_{17}$* | $7.8x10^{-12}$ | **0.23** | 0.0025 | **0.017** | $4.2\ x10^{-06}$ | - |
| *P-value vs $GRS_{228}$* | 0.0077 | **0.35** | **0.82** | **0.22** | **0.2** | - |
| TD | 0.859 | **0.882** | **0.870** | **0.846** | 0.851 | 0.835 |
| *P-value vs $HDQ_{17}$ + $GRS_{228}$* | $1.1\ x10^{-11}$ | 0.0078 | **0.042** | 0.009 | $1.3\ x10^{-10}$ | - |
| *P-value vs $HDQ_{17}$* | $5.8\ x10^{-38}$ | $3.6\ x10^{-07}$ | 0.00024 | $8.8\ x10^{-07}$ | $1\ x10^{-29}$ | - |
| *P-value vs $GRS_{228}$* | $1.6\ x10^{-17}$ | 0.00046 | **0.41** | 0.00024 | $8.1\ x10^{-15}$ | - |
| ROM | 0.803 | 0.798 | 0.835 | 0.803 | 0.805 | 0.791 |
| *P-value vs $HDQ_{17}$ + $GRS_{228}$* | $3.3x10^{-77}$ | $1.5\ x10^{-38}$ | $3.6\ x10^{-6}$ | $3.9\ x10^{-10}$ | $1.7\ x10^{-30}$ | - |
| *P-value vs $HDQ_{17}$* | $2.3\ x10^{-188}$ | $5.4\ x10^{-68}$ | $1.9\ x10^{-14}$ | $6.7\ x10^{-29}$ | $2.5\ x10^{-70}$ | - |
| *P-value vs $GRS_{228}$* | $1.6\ x10^{-122}$ | $4.3\ x10^{-51}$ | $1.5\ x10^{-06}$ | $7.3\ x10^{-21}$ | $6\ x10^{\wedge -45}$ | - |

**Supplementary Table 11:** Frequency, P-values and odds ratios for interactions between common (AF>1%) haplotypes and known CD risk factors for discovery (UK2) and validation (Combined) cohorts. Interactions were only included if the two haplotypes occurred in more than 1% of the UK2 population (Freq). OR indicates the odds ratio and the associated confidence interval. P indicates the –log10(P-value) for the interaction based on a likelihood ratio test. Grey shading is used to separate groups of interactions with the same novel haplotype. Bolding is used to indicate the two interactions that are significant in the discovery and replication cohorts. The threshold for significance here is 0.05/15=0.0033

| | Novel Haplotype | | | Known Haplotype | UK2 | | | Combined | | |
|---|---|---|---|---|---|---|---|---|---|---|
| $G_{NE}$ | HLA-DQA1 | HLA-DQB1 | Symbol | Symbol | Freq | OR (95% CI) | P | Freq | OR (95% CI) | P |
| **1** | **01.02** | **06.02** | **DQ6.2** | **DQ2.5** | **6.0%** | **4.11 (2.55-6.64)** | **4.7E-09** | **6.0%** | **2.53 (1.56-4.13)** | **7.4E-05** |
| 2 | 01.02 | 06.02 | DQ6.2 | DQ2.2 | 2.0% | 0.24 (0.07-0.83) | 3.5E-02 | 1.0% | 0.36 (0.12-1.05) | 8.9E-02 |
| 3 | 01.02 | 06.02 | DQ6.2 | DQ8 | 2.0% | 0.27 (0.12-0.63) | 3.6E-03 | 2.0% | 0.33 (0.15-0.71) | 1.9E-01 |
| 4 | 01.02 | 06.02 | DQ6.2 | DQ7 | 2.0% | 0.48 (0.15-1.51) | 4.2E-01 | 2.0% | 1.19 (0.42-3.41) | 3.9E-04 |
| 5 | 01.01 | 05.01 | | DQ2.5 | 4.0% | 0.74 (0.46-1.19) | 4.8E-01 | 5.0% | 2.12 (1.31-3.43) | 3.7E-04 |
| 6 | 01.01 | 05.01 | | DQ2.2 | 2.0% | 0.2 (0.03-1.53) | 1.4E-01 | 2.0% | 0.47 (0.19-1.15) | 7.1E-02 |
| 7 | 01.01 | 05.01 | | DQ8 | 2.0% | 1.46 (0.69-3.09) | 6.2E-01 | 2.0% | 0.73 (0.38-1.41) | 3.9E-02 |
| 8 | 01.01 | 05.01 | | DQ7 | 2.0% | 1.09 (0.34-3.46) | 1.0E+00 | 2.0% | - | - |
| 9 | 01.03 | 06.03 | | DQ2.5 | 2.0% | 0.73 (0.39-1.36) | 6.2E-01 | 2.0% | 0.94 (0.54-1.65) | 2.2E-02 |
| **10** | **03.03** | **03.01** | **DQ7.3** | **DQ2.5** | **2.0%** | **0.13 (0.06-0.27)** | **1.7E-07** | **1.0%** | **0.19 (0.09-0.42)** | **3.8E-01** |
| 11 | 03.03 | 03.01 | DQ7.3 | DQ2.2 | 1.0% | 1.69 (0.37-7.69) | 8.1E-01 | 1.0% | 2.1 (0.46-9.54) | 2.9E-10 |
| 12 | 03.03 | 03.01 | DQ7.3 | DQ8 | 1.0% | 5.19 (2.18-12.32) | 3.2E-03 | 1.0% | 3.33 (1.25-8.87) | 1.3E-09 |
| 13 | 03.03 | 03.01 | DQ7.3 | DQ7 | 1.0% | 3.06 (0.82-11.39) | 3.2E-01 | 1.0% | 14.35 (4.25-48.53) | 3.0E-49 |
| 14 | 01.02 | 06.04 | | DQ2.5 | 1.0% | 1.03 (0.49-2.18) | 1.0E+00 | 1.0% | 1.19 (0.6-2.36) | 4.4E-03 |
| 15 | 02.01 | 03.03 | | DQ2.5 | 1.0% | 0.24 (0.09-0.61) | 1.7E-02 | 1.0% | 1 (0.37-2.71) | 1.9E-03 |

**Supplementary Table 12: Impact of shifting exclusionary criteria if we consider 100,000 tests with prevalence of either 1% or 10%.** The numbers below are based on the average positive and negative predictive values derived from each of the five cohorts in our study, as shown in Figure 4 in the main text. The exclusionary criteria (i.e. genotypes that indicate a negative CD diagnosis) are as follows: current is DQX/DQX, DQ7/DQX, DQ7/DQ7, current+1 adds DQ8/DQ7, current+2 adds DQ2.2/DQX, current+3 adds DQ8/DQX, as per Figure 4 in the main document.

| Prev | Exclusion criteria | TP | FP | TN | FN | PPV | NPV |
|---|---|---|---|---|---|---|---|
| 1% | DQX/DQX DQ7/DQX, DQ7/DQ7 | 982 | 51667 | 47333 | 18 | 0.019 | 1.000 |
| | DQX/DQX, DQ7/DQX, DQ7/DQ7, DQ8/DQ7 | 977 | 49512 | 49488 | 23 | 0.019 | 1.000 |
| | DQX/DQX, DQ7/DQX, DQ7/DQ7, DQ8/DQ7, DQ2.2/DQX | 962 | 40045 | 58955 | 38 | 0.023 | 0.999 |
| | DQX/DQX, DQ7/DQX, DQ7/DQ7, DQ8/DQ7, DQ2.2/DQX, DQ8/DQX | 935 | 28293 | 70707 | 65 | 0.032 | 0.999 |
| 10% | DQX/DQX DQ7/DQX, DQ7/DQ7 | 9818 | 46970 | 43030 | 182 | 0.173 | 0.996 |
| | DQX/DQX, DQ7/DQX, DQ7/DQ7, DQ8/DQ7 | 9769 | 45011 | 44989 | 231 | 0.178 | 0.995 |
| | DQX/DQX, DQ7/DQX, DQ7/DQ7, DQ8/DQ7, DQ2.2/DQX | 9619 | 36404 | 53596 | 381 | 0.209 | 0.993 |
| | DQX/DQX, DQ7/DQX, DQ7/DQ7, DQ8/DQ7, DQ2.2/DQX, DQ8/DQX | 9354 | 25721 | 64279 | 646 | 0.267 | 0.990 |

### Supplementary Figures

**Supplementary Figure 1:** ROC curves of all models considered in this paper across the FIN, IT, NL and UK1 cohorts. Each corresponds to a different model and the corresponding AUC is shown in the legend. Staring of the legend entry indicates that the model is statistically equivalent to the best performing model.

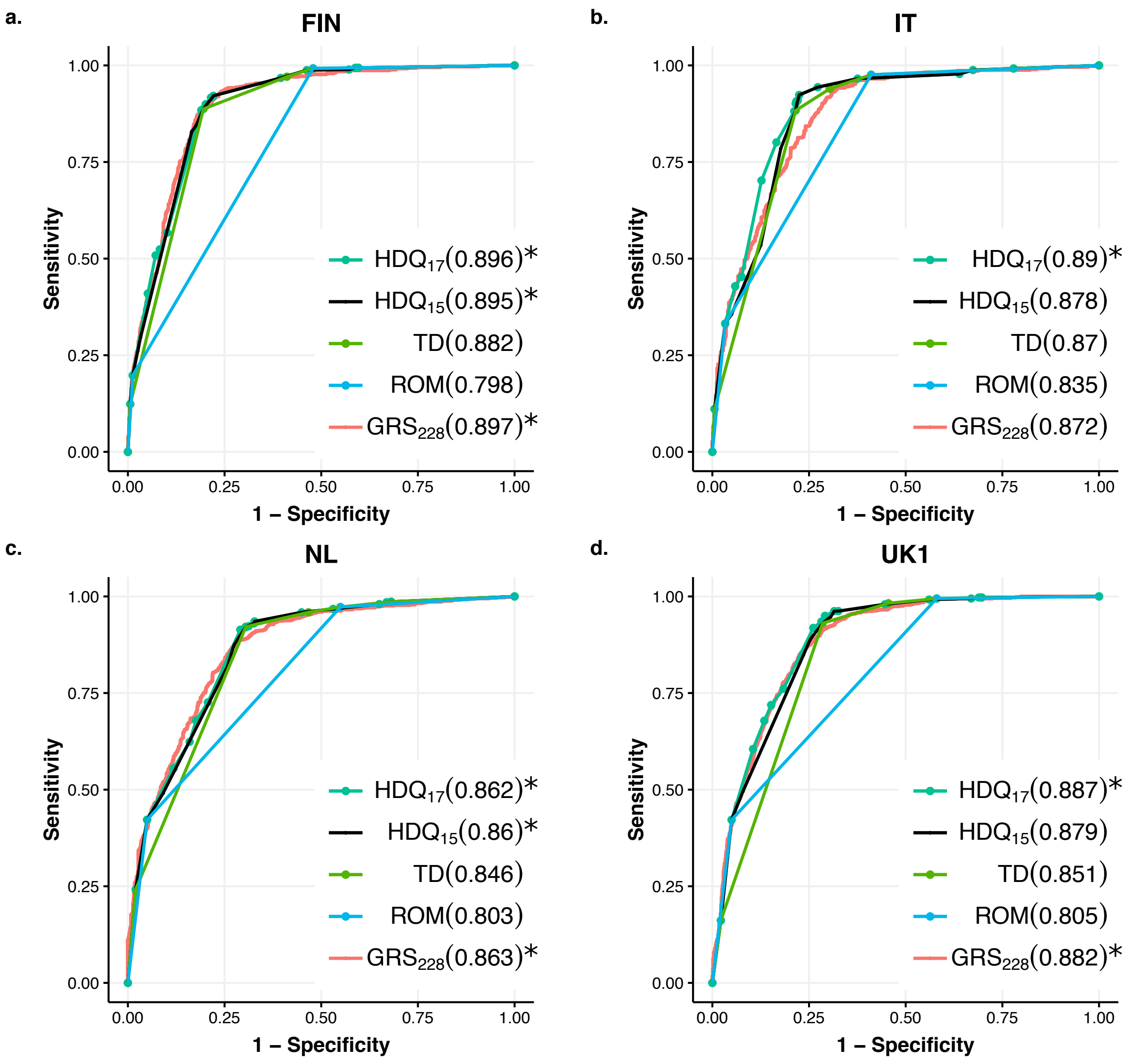

**Supplementary Figure 2: Performance of proposed HLA models using either imputed haplotypes or 4-6 tag SNPs.** Left and right subplots represent the $HDQ_{15}$ and $HDQ_{17}$ models respectively, with the set of points indicating AUCs if the model were based on imputed haplotypes (Hap.) or tag SNPs (SNP), 4 SNPs for $HDQ_{15}$ and 6 for $HDQ_{17}$. No significant differences were found between the use of imputed haplotypes or SNPs for either model as evaluated by DeLong's test. Horizontal lines represent the mean AUC.

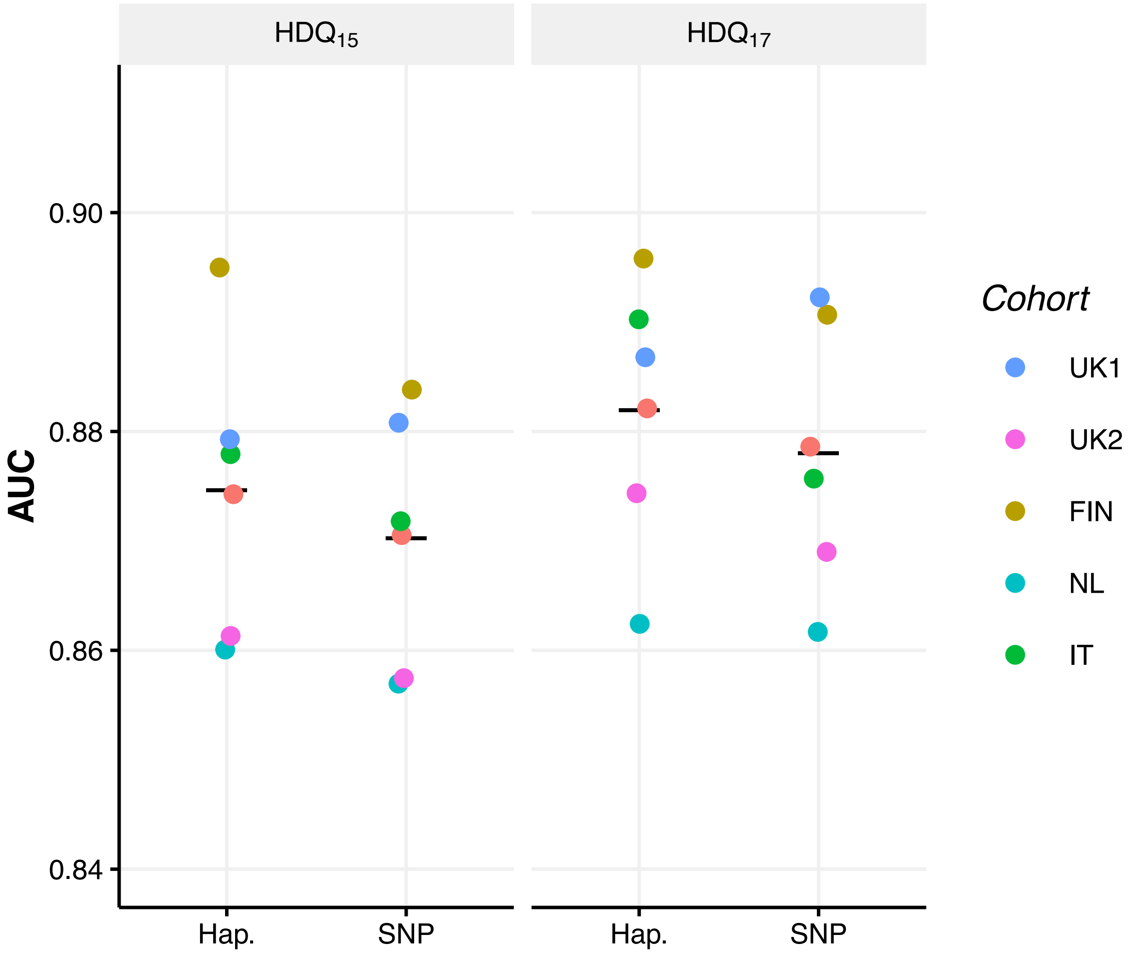

**Supplementary Figure 3**

To assess whether predictive performance could be further improved through combination of the $HDQ_{17}$ and $GRS_{228}$ models, the performance of models variably weighting each predictor were assessed. No significant improvement could be observed over either model using this combination approach. The combination presented in the paper makes use of a 0.9 weighting of HDQ17, meaning the resulting combination reorders the samples within each of the 17 $HDQ_{17}$ categories by the $GRS_{228.}$

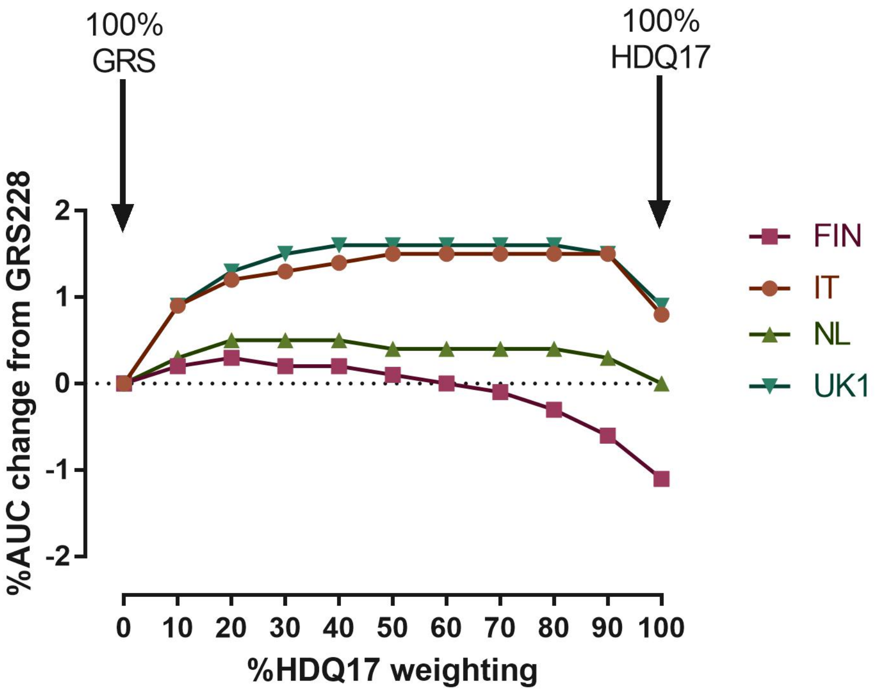
